# A Pipeline for Screening Small Molecule-Enhanced Protein Stability in A Bacterial Orphan Receptor

**DOI:** 10.1101/2025.07.03.663076

**Authors:** James J. Siclari, Denize C. Favaro, Richard H. Huang, Kevin H. Gardner

## Abstract

Bacterial one-component signaling proteins integrate sensory and gene regulation functions within the same polypeptide, creating powerful natural sensors of environmental conditions which can also be adapted into powerful tools for synthetic biology and biotechnology. A key sensor motif within many of these proteins is the Per-ARNT-Sim (PAS) domain, known for its conserved fold yet highly divergent sequences, allowing for a broad range of ligands to control PAS protein function by changes in small molecule binding occupancy or configuration. This diversity introduces a challenging step – identification of ligands for “orphan” PAS proteins which show signatures of ligand binding but no copurifying high-affinity bound small molecules – into characterization and engineering of such proteins. In this study, we characterized CU228, a putative PAS-HTH transcription factor from *Candidatus Solibacter usitatus,* as a novel model system for searching for novel ligands by small molecule stabilization. Bioinformatics and structural analyses predicted a PAS domain with an Trp-rich internal cavity, suggesting potential small molecule interactions. Using a ∼760 compound fragment library, differential scanning fluorimetry identified three ligands (KG-96, KG- 408, and KG-484) that substantially increased CU228 thermal stability with ΔT_m_ values up to +10°C. Microfluidic modulation spectroscopy (MMS) revealed ligand-induced preservation of α-helical and β-sheet integrity under thermal stress. Saturation transfer difference NMR confirmed direct binding of all three ligands and enabled estimation of micromolar-range dissociation constants, consistent with expected fragment-level affinity. Our findings expand the analytical toolbox for probing protein-ligand interactions in flexible, signal-responsive systems, laying the groundwork for designing synthetic chemogenetic variants of one-component transcription factors.

## Introduction

Responding to an ever-changing external environment is crucial for bacterial survival and proliferation. Many such adaptive responses are transcription factors based on one component signaling proteins (Ulrich et al. 2005), which couple a sensor domain and a DNA-binding domain within the same polypeptide chain, switching gene expression on and off in response to changes in ligands from their surroundings.

Exploiting the natural functions of bacterial proteins has driven significant advancements in biotechnology; indeed, natural one component signaling proteins have laid the foundations for many biosensors and, inducible expression systems, often through slight modifications to existing functionalities (Lazar and Tabor 2021).

One highly used sensor domain in many of these proteins is the Per-ARNT-Sim (PAS) domain, which is characterized by a highly conserved ∼120 amino acid residue mixed ɑ/β fold and often containing an internal hydrophobic pocket evolved for ligand binding. While the structural fold is highly conserved, sequence identity across PAS domains is less than 20%, enabling them to sense a wide range of stimuli (Taylor and Zhulin 1999; Henry and Crosson 2011; Stuffle et al. 2021).

An example of this from our lab is the blue-light-sensitive PAS transcription factor EL222, which contains an N-terminal light-oxygen-voltage LOV domain (a subfamily of PAS domains, evolved to bind flavin chromophores) and a C-terminal DNA binding helix-turn-helix (HTH) domain. EL222 undergoes a conformational change upon blue light exposure, transitioning from a monomer to a homodimer and enabling DNA binding (Nash et al. 2011; Rivera-Cancel et al. 2012; Zoltowski et al. 2013; Chaudhari et al. 2025). Previous work has engineered this system for multiple uses in eukaryotic model systems (Motta-Mena et al. 2014; Reade et al. 2016; Rullan et al. 2018; Zhao et al. 2018; Zhao et al. 2020; Cleere and Gardner 2024; Hoffman et al. 2025). LOV domains are easily identifiable by their primary sequences, including a conserved “NCRFL” motif that is used to bind flavin chromophores like FMN, FAD, and riboflavin (Henry and Crosson 2011). While many other PAS domain subfamilies exist in one-component systems and likely respond to a wide variety of ligands, the sequence conservation and ligand-binding specificities of these domains is less well understood.

Identifying and engineering LOV domains has the added advantage that LOV- binding cofactors are naturally available in organisms typically used for heterologous expression and bind with sufficient affinity for complexes to survive *in vitro* purification and characterization. Because this is not always the case for all ligands, we have built an in-house library of approximately 760 small-molecule fragments that has been useful for ^15^N/^1^H HSQC-based NMR-based screening of several PAS proteins (Amezcua et al. 2002; Scheuermann et al. 2009; Guo et al. 2013; Xu et al. 2021) and non-PAS proteins (Best et al. 2004). While this strategy has been successful for identifying useful compounds – some of which led to the successful development of a FDA-approved PAS domain inhibitor targeted cancer therapeutic (Cho et al. 2016; Jonasch et al. 2021), such protein-detected NMR screening approaches can be laborious due to high sample consumption.

In this study, we aimed to use this library to identify small molecule ligands which could bind and potentially activate a novel PAS-HTH protein as a new chemogenetically-controlled transcription factor. Unfortunately, the PAS-HTH protein we selected here, CU228, was not well behaved for protein-directed NMR screening, requiring us to develop a novel multifaceted screening pipeline incorporating differential scanning fluorimetry (DSF), ligand-directed nuclear magnetic resonance (NMR), and microfluidic modulation spectroscopy (MMS). From this approach, we identified compounds that clearly bound CU228 with accompanying changes in protein stability suggestive of conformational changes within this novel PAS-HTH protein. Our results both seed the potential development of novel CU228-binding ligands and lay out a general strategy for rapidly conducting both binding and stabilization assays to identify new compounds capable of binding moderately behaved targets.

## Results

### Identification of a Protein Test Case

To identify a suitable test case for small molecule stabilization, we searched the SMART database (Letunic et al. 2021) for predicted one-component prokaryotic proteins containing both PAS and HTH domains. From this list of ∼2600 sequences, we subsequently used a bioinformatics pipeline to select sequences with a single PAS domain (lacking the canonical NCRFL flavin-binding motif) and flanked to either side by a single HTH domain (**Fig. 1a**) (Edgar 2004; Cai et al. 2004; Robert and Gouet 2014; Zimmermann et al. 2018; Gabler et al. 2020; Gumerov and Zhulin 2020; Letunic et al. 2021). Out of the fifty sequences remaining at the end of this pipeline, we focused on a 228-residue protein from *Candidatus Solibacter usitatus* (Challacombe et al. 2011) (CU228). Conspicuously, AlphaFold3 modeling (Abramson et al. 2024) predicted that three tryptophan residues within the PAS domain would arrange to form a cage-like structure that could bind an internal ligand (**Fig. 1b**). We assessed the structural integrity, secondary structure composition, and oligomeric state of CU228 using microfluidic modulation spectroscopy (MMS, an IR-based spectroscopy focused on the amide-I region) (Liu et al. 2020) and size exclusion chromatography coupled to multi- angle light scattering (SEC-MALS). Taken together, these studies revealed that the protein was a stable dimer and contained a mixed α/β fold with a particularly strong helical content, consistent with PAS-HTH architecture (**Fig. 1c,d**). Complementary ^15^N/^1^H TROSY NMR experiments (**Fig. S2**) displayed poor amide ^1^H chemical shift dispersion and heterogeneous peak intensities, suggesting heterogeneous tertiary structure. Taken together, our data indicated CU228 is a structured yet dynamic and flexible protein, making it an attractive candidate for stabilization through small-molecule binding.

**Figure 1.**
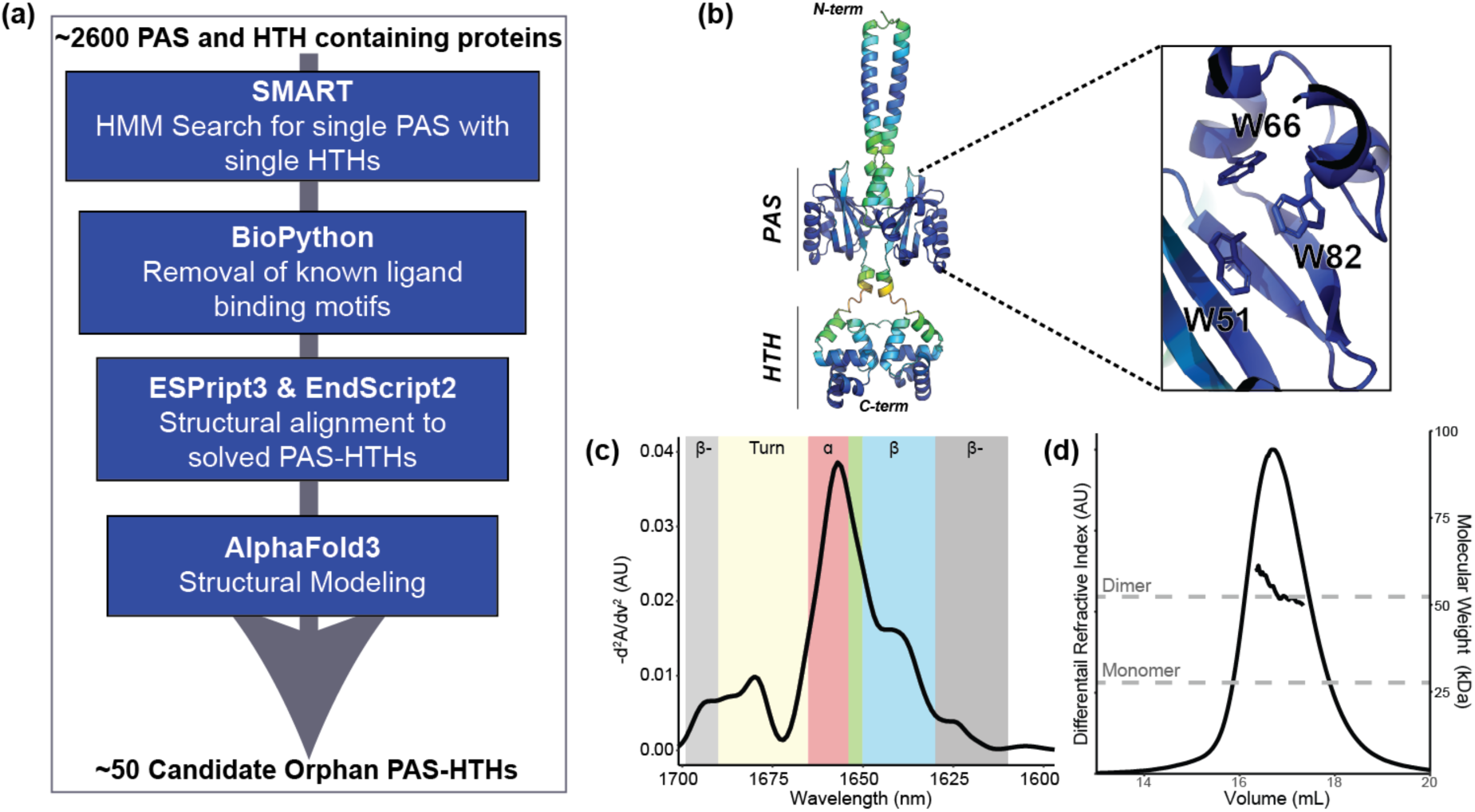
Initial characterization of test case protein for this study, CU228. (a) Schematic of bioinformatics pipeline for combinatorial domain hunting searching for a PAS-HTH with no FMN binding preferences. (b) AlphaFold3 model colored by pLDDT score (blue = high confidence to red = low confidence), with zoom inset showing three internally facing tryptophan side chains in the PAS domain cavity. (c) MMS spectra of purified CU228 at 25°C, colors denote secondary structure elements, can be seen in Table S1. β-: Intermolecular β-sheet, α: α-helix, β: β-sheet, green box: unordered structure. (d) Superdex 200 SEC-MALS chromatogram of CU228. Vertical axes indicate relative protein concentration via differential refractive index (dRI, arbitrary units) and MALS-derived mass is shown as a solid line under the peak. Dashed grey lines indicate the calculated monomer and dimer masses based on primary sequence.

### Screening for Small Molecule Stabilizers

We conducted a thermal shift assay using DSF to identify ligands that enhanced CU228 stability (**Fig. 2**). To capture medium-affinity binders, we screened our 760 compound fragment library (Amezcua et al. 2002) with 2.5 µM CU228 and 250 µM of each small molecule, each with 1.25% DMSO and performed in duplicate. CU228 alone exhibited a melting temperature (T_m_) of 53.4±0.07°C in the presence of 1.25% DMSO, which we confirmed to have minimal effect on the CU228 structure by 1D ^1^H NMR (**Fig. S3**). While the T_m_ of CU228 was minimally impacted by the addition of most compounds in the library, approximately 50 compounds stabilized CU228 at least one standard deviation above the control (T_m_ > 55.7°C) (**Fig. 2a**). At this juncture, we discarded CU228/ligand complexes with poor reproducibility or solubility (suggesting practical complications with further biophysical analysis) or biphasic melting curves (suggesting more complex protein:compound:dye interactions than ideal for further optimization).

**Figure 2.**
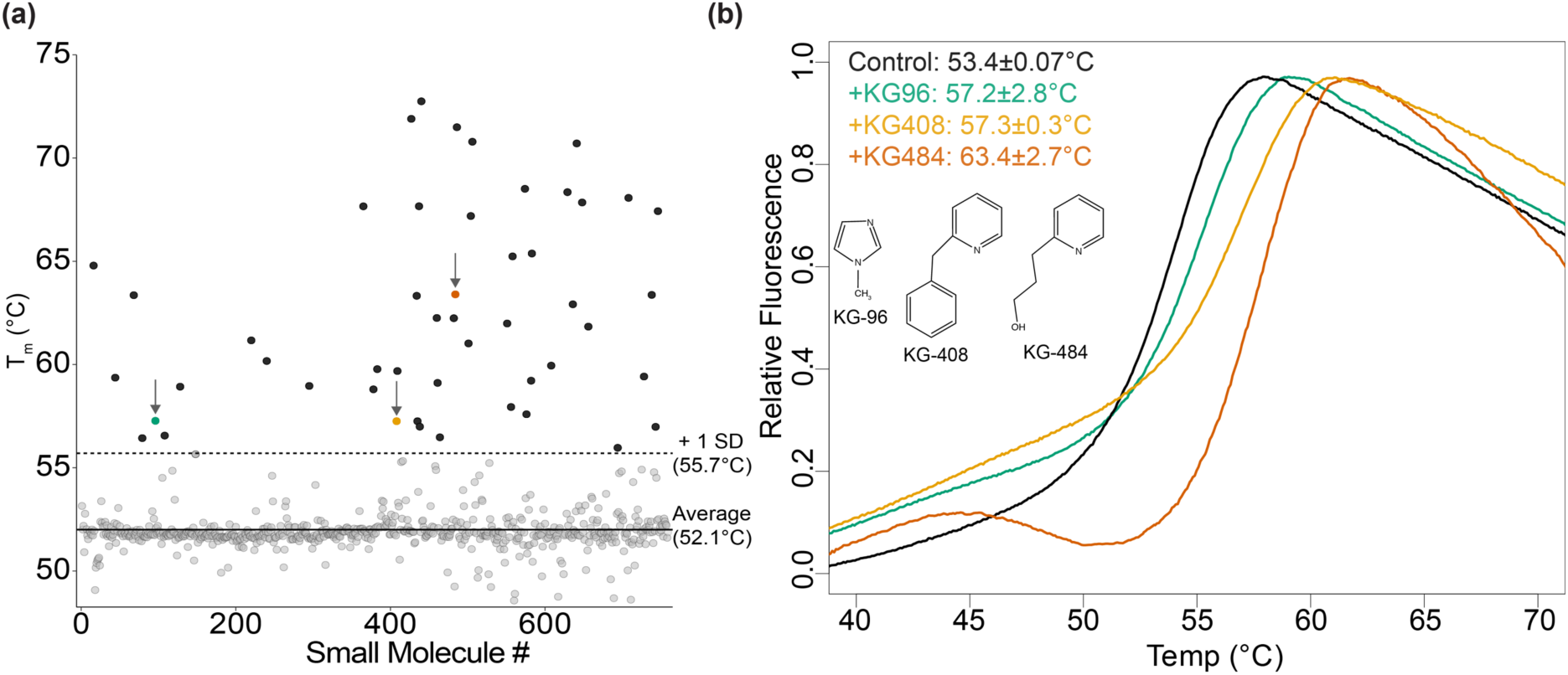
Initial CU228 small molecule fragment screen by thermal shift assay. (a) Average T_m_ (n=2) for all compounds in the screen calculated by thermal shift. (b) Raw fluorescence traces of Protein Thermal Shift^TM^ Dye associating and dissociating to CU228 as a function of temperature in the control sample (T_m_ = 53.4±0.07°C) and three ligands: green: KG-96 (T_m_ = 57.2±2.8°C), yellow: KG-408 (T_m_ = 57.3±0.3°C), and red: KG-484 (T_m_ = 63.4±2.7°C). Insert shows chemical structure for respective compounds.

This left us with three ligands for further characterization: KG-96 (T_m_ = 57.2±2.8°C), KG- 408 (T_m_ = 57.3±0.3°C), and KG-484 (T_m_ = 63.4±2.7°C). Notably, all three hit compounds included nitrogen-containing heterocycles and small aliphatic or aromatic side chains. (**Fig. 2b**).

### Validation of Ligand Binding and Affinity Determination via NMR

Given the low quality of the apo CU228 ^15^N/^1^H-TROSY spectrum (**Fig. S2**), we wanted to establish if we could obtain higher quality spectra of CU228 in the presence of any of the three compounds identified by thermal shift assay. Unfortunately, while we did not see any substantial improvement in TROSY peak dispersion or linewidths by the addition of 1 mM KG-96, KG-408, or KG-484, we saw ligand-dependent changes at several specific sites throughout the spectra (**Fig. S4a**, red arrows). While were unable to get sequence-specific assignments of these spectra to get precise locations, we suggest peaks in two conspicuous spectral regions have known locations in CU228.

One of these, with very downfield ^1^H shifts ∼11 ppm (**Fig. S4b**), corresponds to an extremely unusual backbone amide environment at the N-terminus of the Fα helix in many other PAS domains (Holdeman and Gardner 2001; Card et al. 2005; Corrêa et al. 2016). We observed a weak apo protein peak in this region, and saw slight ligand-dependent chemical shift changes by addition of either KG-96 or KG-484, providing some validation of both the PAS domain being present and involved in ligand binding. In a second region with downfield ^1^H and ^15^N shifts (**Fig. S4c**), where tryptophan indole sidechain NH signals commonly appear, we saw one of the two sets of observed peaks show substantial ligand-dependent chemical shift changes. Given that all four Trp residues of CU228 are within the PAS domain, this provides strong evidence of chemically-triggered structural changes involving this predicted ligand-binding domain.

To confirm the direct binding of the three ligands to CU228, we used saturation transfer difference (STD) NMR, a ligand-observed NMR method where protein (10-50 µM) was incubated in the presence of increasing concentrations of ligand (0.02-5 mM). We acquired STD NMR data with an interleaved acquisition of two spectra, one with protein signals saturated (ON) and one without (OFF). Differences in intensity between these ON and OFF spectra reveal a transfer of saturation via intermolecular Nuclear Overhauser Effect (NOE) (Angulo et al. 2010) from a protein/ligand complex, indicating direct binding. We observed clear ligand peaks in the STD difference spectra, indicating saturation transfers from CU228 to KG-96, KG-408, and KG-484 consistent with direct interactions in the fast exchange regime (**Fig. 3a**).

**Figure 3.**
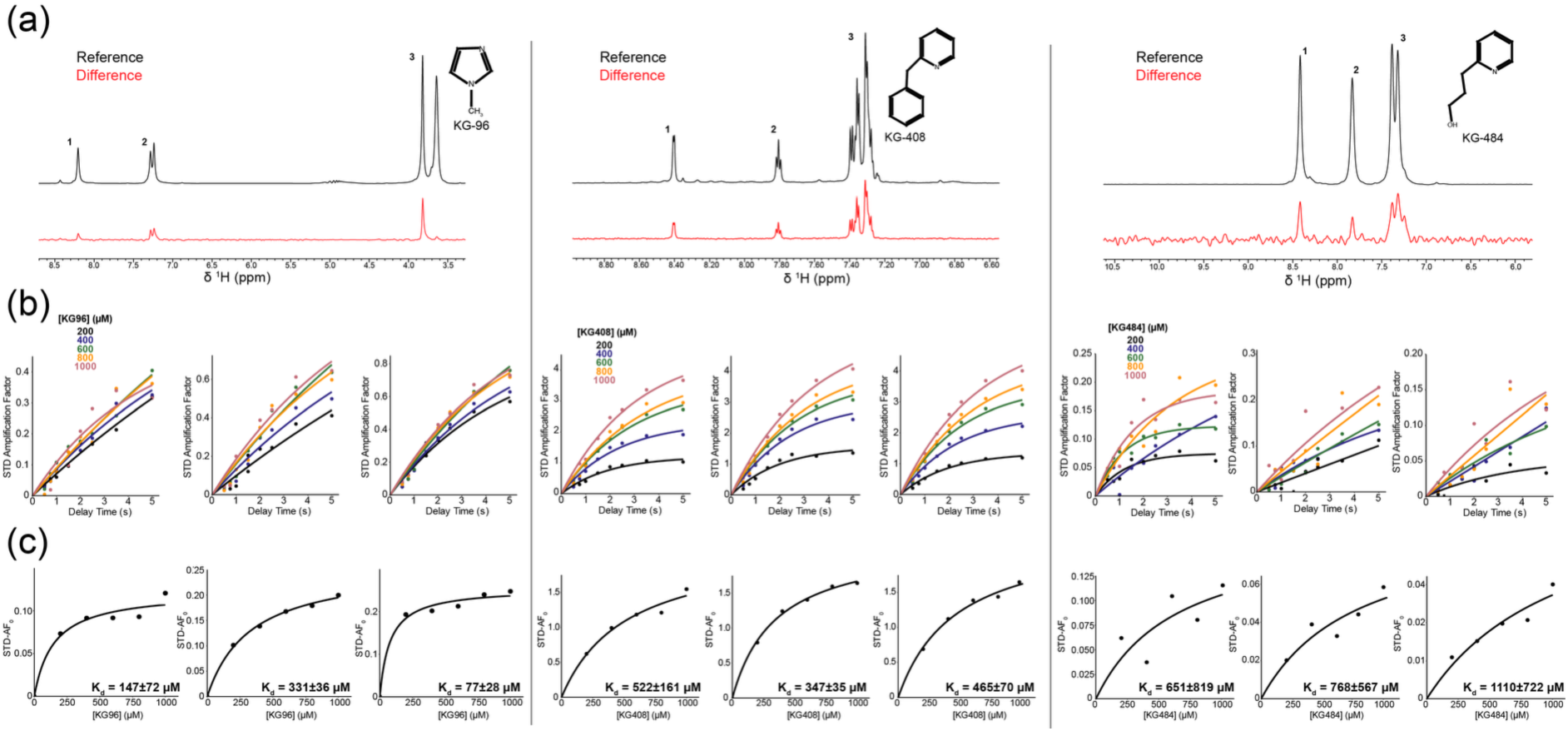
**STD NMR of CU228 and KG-96, KG-408, or KG-484**. (a) ^1^H spectra of reference (black) and difference (red) for STD experiments (t_sat_ = 2 s) between CU228 and the respective compound at 1 mM, and 30 µM CU228. The insert shows the chemical structure of each. (b) STD buildup curves for each spectral region labeled in panel a utilizing varying saturation times. (c) STD binding curves for each spectral region labeled in panel a. Weighted average K_d_ from three epitopes for KG- 96, KG-408, or KG-484 were 171±21 µM, 376±31 µM, and 841±392 µM respectively.

To determine binding affinities, STD experiments were performed at various ligand concentrations and saturation times. We calculated STD amplification factors (AFs) for three spectral regions for each ligand (**Fig. 3b**; **Eq. 1-2**). We then generated binding curves by plotting amplification factor growth rates (STF-AF_0_) versus ligand concentration and fitting each to a Langmuir isotherm (**Eq. 3**), yielding average dissociation constants (K_d_) for KG-96, KG-408, and KG-484 of 171±21 µM, 376±31 µM, and 841±392 µM respectively (**Fig. 3c**) (Mayer and Meyer 2001; Angulo et al. 2010).

### Assessing Secondary Structure Changes via MMS

While the thermal shift assay works well to probe overall tertiary structure, it does so by testing the accessibility of a dye to the protein hydrophobic core, raising potential concerns in the case of suspected small molecule binders such as CU228 and more generally (Wu et al. 2023). To further assess the structural impacts of ligand binding on CU228 without any bias or competition from a dye molecule, we employed MMS to monitor secondary structure changes of CU228 in the presence of KG-96, KG-408, and KG-484.

At ambient temperature, each ligand had minimal effects on the CU228 secondary structure, aside from KG-408’s ability to increase both native and intermolecular β-sheet content (β-) (**Fig. S5**). Upon exposure to thermal stress, however, all three ligands stabilized β-sheet and α-helix at elevated temperatures, relative to the control as indicated by the increased T_m_ values (**Fig. 4, Table S2**). Additionally, ligand-treated samples exhibited delayed formation of β-aggregates during heating (**Fig. 4**), suggesting ligand-protein interactions might stabilize the dimeric structure involving predicted PAS domain β-sheets in the CU228 AlphaFold3 model.

**Figure 4.**
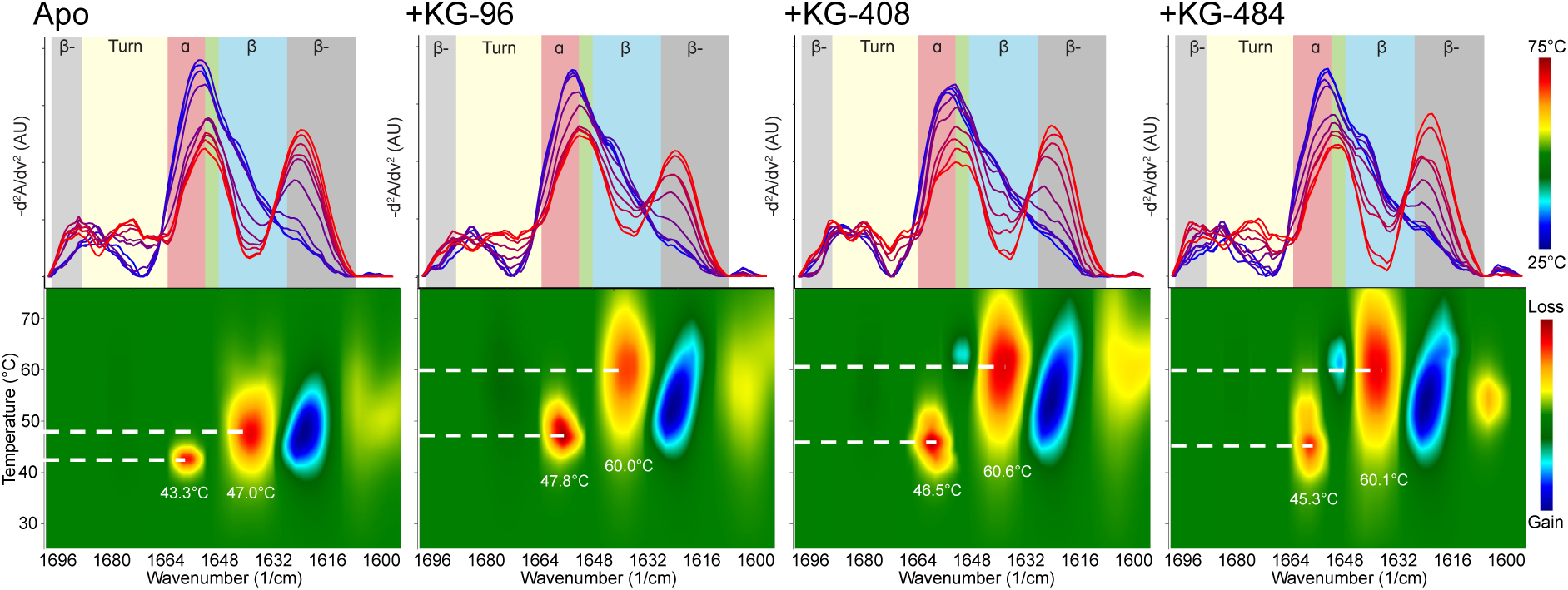
Thermal ramp MMS of apo and ligand-bound CU228. *(top row)* IR spectra of 20 µM CU228, either alone or in the presence of 1 mM of indicated ligand, at 5°C intervals between 25-75°C, displaying changes in aggregated β-sheet (grey), turn (yellow), ⍺-helices (red), unorder (green), and native β-sheet (blue). *(bottom row)* Heat maps displaying temperature-dependence of IR signal change in CU228. Blue denotes increase in spectral intensity or gain in respective structure; red denotes decrease in spectral intensity or loss in respective structure. Dotted lines point to temperature with the highest rate of range, indicative of T_m_.

The degree of thermal stability was concentration dependent, with ranges comparable to K_d_ measurements acquired via NMR (**Fig. 5**). Maximal fractional contribution for α- helical and β-sheet content arises at temperatures around 35-40°C, which was not seen in the apo protein (**Fig. 5**), suggesting that full folding of CU228 requires the presence of a small molecule ligand.

**Figure 5.**
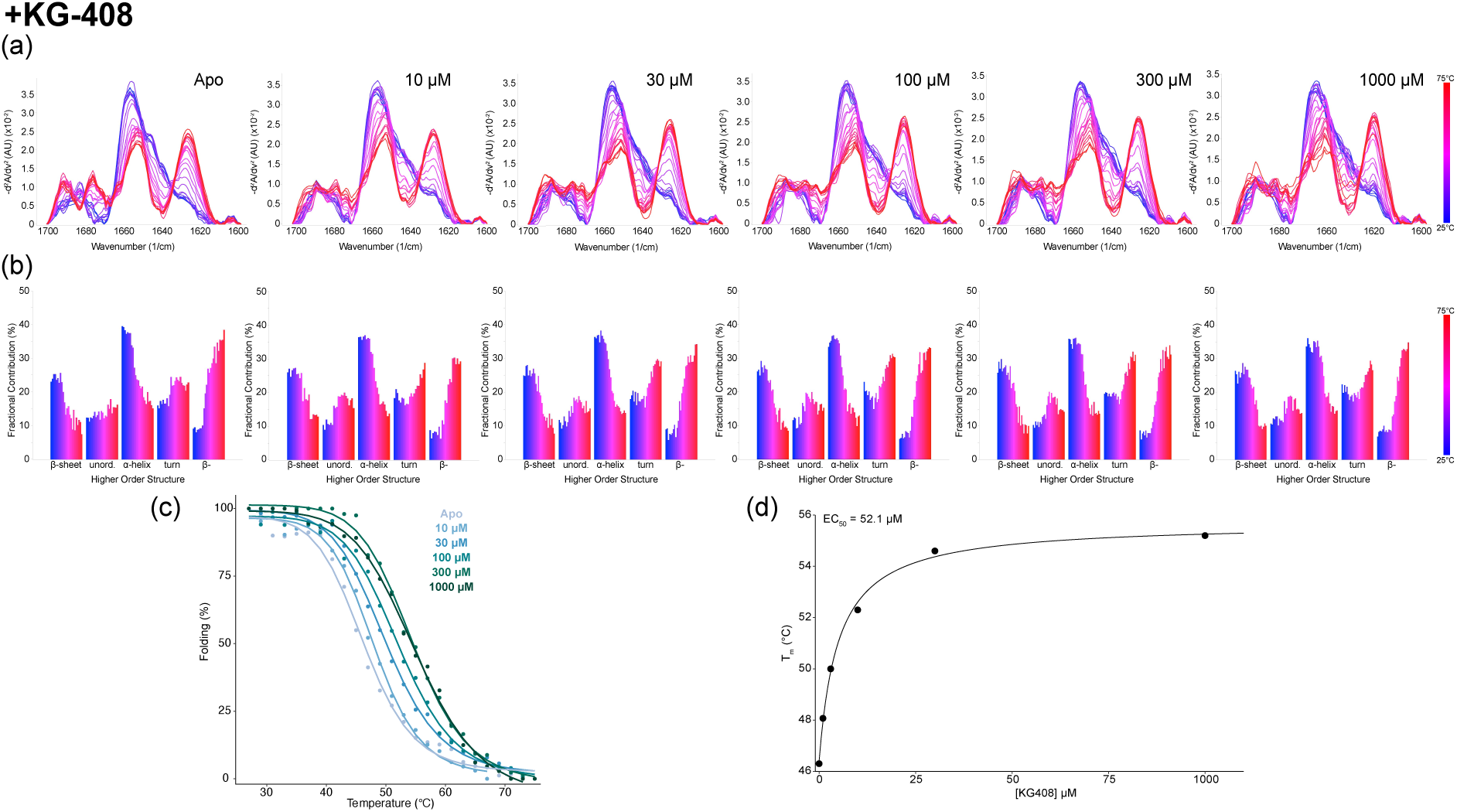
Concentration dependent thermal ramp MMS of CU228 with KG-408. (a) MMS similarity spectra across from 25-75°C at five different concentrations of KG-408. (b) Higher order structure fractional contribution as a function of temperature for each titration point. (c) Global melting curves of CU228 with addition of each KG- 408 concentration. (d) KG-408 impact on the melting temperature of CU288 as a function of ligand concentration.

temperatures of all secondary structures concomitantly increased (**Fig. 5b)**. However, while α-helical T_m_ values were moderately affected (4-5°C) by ligand binding, β-sheet resistance to thermal denaturation was increased more substantially, up to 14°C (**Table S2**). We interpret this observation as supporting evidence of the binding of these small compounds within the mixed α/β fold of the CU228 PAS domains, as the only predicted beta-sheet containing element in the protein. In contrast, the interdomain linker and DNA binding domain of the protein are predicted to be all helical, and are accordingly less stabilized by ligand binding. Globally, the T_m_ CU228 increased in a concentration dependent manner (**Fig. 5c**). Each T_m_ was then plotted to estimate an effective concentration (EC_50_) of ligand-driven thermal stability (**Fig. 5d**). While **Fig. 5** presents data from KG-408, the two other ligands KG-96 and KG-484 showed similar results (**Fig. S6, S7**).

We also noticed changes to the 1600 nm region of the IR spectra during thermal stress, which is typically attributed to changes in amino acid side chains, particularly Tyr (Kong and Yu 2007; De Meutter and Goormaghtigh 2021). This was mainly observed with CU228 binding to KG-96 (**Fig. 4**), which coincidentally is the smallest of the three ligands tested, perhaps further implicating interior binding near the two tyrosine residues located within the PAS core.

## Discussion

This study presents a multifaceted workflow for identifying and characterizing ligand interactions with a previously unstudied orphan PAS-HTH transcription factor. A structure-guided bioinformatic approach identified CU228 as a promising candidate, with a predicted Trp-rich pocket likely suitable for small-molecule targeting. Through thermal shift assays, multiple fragments were found to increase protein thermal stability. MMS provided insights into the preservation of secondary structure under thermal stress, and NMR confirmed direct ligand binding while quantitating high-micromolar binding affinities.

Historically, protein receptors have been deorphanized *in vitro* through mass spectrometry or atomic structure determination to identify the natural ligands (Laschet et al. 2018). This approach requires the ligand to be present in the expression system and be of high enough affinity to remain bound through all sample preparation steps.

Screening of artificial ligands has been beneficial in guiding natural ligand discovery as well as developing triggers for protein engineering but can require a large amount of sample when run in certain formats. For example, our prior ^15^N/^1^H HSQC NMR-based screening of our 760 compound library against PAS domains (Amezcua et al. 2002; Scheuermann et al. 2009; Guo et al. 2013) or other small proteins (Best et al. 2004) typically entailed the use of 150-200 mg of uniformly ^15^N-labeled protein. While this protein-detected NMR approach is extremely data-rich, providing very early binding site information and affinity, the sample requirements limit its application to many systems. In contrast, the pipeline described here requires a low quantity of protein – approximately 4 mg for all of the experimental work describe herein – and can be scaled up to perform high-throughput screens.

The relatively low affinities we observed for each ligand is consistent with typical expectations for fragment-based screening, where focused collections of small compounds (MW ∼ 200 Da) are used to quickly screen a fraction of chemical space to lay the foundation for further optimizations (Barelier and Krimm 2011; Kirkman et al. 2024). More specifically, the mid- to high micromolar affinities seen for KG-96, KG-408, and KG-484 are also consistent with our experience with this small fragment library against other targets with affinities between 100-500 µM (Amezcua et al. 2002; Best et al. 2004; Guo et al. 2013), some of which were rapidly improved to low micromolar or high nanomolar potency (Scheuermann et al. 2009; Key et al. 2009). In addition to getting these initial leads, we also obtained some early structure-activity relationship (SAR) information that could assist future development of higher affinity derivatives from these fragment scaffolds. The potential for this can already be seen as the difference of a phenyl ring between KG-408 and KG-484 results in the doubling of K_d_ value.

In addition to developing a deorphanization strategy, our findings offer new insights into the confirmational behavior of CU228 that can be extended to other one-component signaling proteins. The observed increase in β-sheet melting temperature upon ligand binding and prolonged intermolecular β-sheet interactions are consistent with the predicted regions for stimulus recognition and homodimerization interfaces within the PAS domain suggest stabilization of the dimer complex though the PAS domain β-sheets. Seeing the greatest structural resilience in the β-sheet content of CU228 further supports that each ligand directly interacts with the PAS domain, while the remaining helical content is less perturbed (**Table S2**).

Accordingly, literature suggests the 1650-1652 cm^-1^ region of the IR spectra is indicative of coiled-coil formation (Heimburg et al. 1999; Dong et al. 2008). Absorption in this region became more prominent in our results when the CU228 was heated above its melting temperature (**Fig. 5**). In the apo state, this prominent peak offers more support for the AlphaFold3 model of CU228 (**Fig. 1**) where the N-terminal ⍺-helices are one of the predicted dimerization interfaces. Upon CU228 binding to KG-96, this coiled-coil region (1650-1652 cm^-1^) retains structure despite thermal stress, while the non- coiled coil α-helical structures (1656 cm^-1^) found in the PAS, linker, and HTH segments melt (**Fig. S6**). These findings are consistent with literature precedence from other dimeric PAS proteins, where ligand-dependent changes within the PAS domain impact the stability and function of flanking helical elements and output domains (Ralph et al. 2013).

Overall, our integrative approach demonstrates the utility of combining computation modeling, biophysical assays, and NMR to study flexible, dynamic proteins and their interactions with small molecules, particularly when no prior knowledge of endogenous ligand exists. Strengths of this approach are its speed, low sample consumption, and mix of label-free and label-containing approaches to quickly get a slate of fragment and protein information for subsequent higher-content assays and optimization, enabling larger classes of protein targets to be surveyed.

## Materials and Methods

### Cloning, Protein Expression and Purification

DNA encoding the CU228 sequence (GenBank: HEY1493056.1) was ordered from Twist Biosciences and cloned into a pHisGβ1-parallel expression vector (Harper et al. 2003). CU228 was recombinantly expressed with a His_6_-Gβ1 tag in BL21(DE3) *E. coli* (Stratagene) in LB media at 16°C overnight. Cells were pelleted and resuspended in Buffer A (50 mM Tris-HCl pH 8.0, 500 mM NaCl). The cells were lysed by sonification, and the lysate was cleared by centrifuging at 10,000 x g for 30 minutes at 4°C. The supernatant was collected and filtered through a 0.2 µm syringe filter.

For affinity purification, the protein was loaded onto a 5 mL HisTrap (Cytiva) column preequilibrated with Buffer A. The column was washed with 10 column volumes (CVs) of Buffer A before eluting the protein with buffer B (50mM Tris-HCl pH 8.0, 500 mM NaCl, 500 mM imidazole). The eluent was diluted 1:10 with Buffer C (50 mM Tris-HCl, pH 8.0) and the His_6_-Gβ1 tag was cleaved by adding 1 mg His_6_-TEV protease (Kapust et al. 2001; Phan et al. 2002) per 30 mg of fusion protein. After overnight cleavage, the sample was again applied to a preequilibrated 5 mL HisTrap column to remove tag and TEV protease. The flow through was then concentrated using an 30 kDa MWCO Amicon Ultra concentrator to a volume of 3 mL. The concentrated flowthrough was applied to a Superdex 200 10/300 (Cytiva) column preequilibrated with assay buffer (10 mM sodium phosphate pH 7.0, 50 mM NaCl, 0.5 mM DTT).

### Size Exclusion Chromatography coupled With Multiangle Light Scattering (SEC-MALS)

All samples and buffers were filtered through a 0.1 µm pore filter before use. CU228 was concentrated to 30 µM and injected onto a Superdex 200 GL 10/300 SEC column pre-equilibrated with assay buffer. Following elution, light scattering was measured by an inline miniDAWN TREOS and refractive index was measured using an Optilab rEX (Wyatt Technology). The entire process was carried out at room temperature. Data analysis and molecular weight calculations were performed using ASTRA V software (Wyatt Technology).

### Thermal Shift Assay

Sourcing for compounds in the small molecule library is outlined in Guo et al., 2013 (Guo et al. 2013). Thermal shift assays were performed using Applied Biosystems Protein Thermal Shift^TM^ Dye kit as per the manufacturer’s instructions. CU228 was used at a concentration of 2.5 µM. All small molecule fragments were diluted to 250 µM in 1.25% DMSO. Fluorescence was recorded using a QuantStudio 7 Flex Real-time qPCR machine. All data were processed and analyzed using TSA-CRAFT (Lee et al. 2019).

### Microfluidic Modulation Spectroscopy

For initial characterization of CU228 and folding confirmation, MMS measurements were made on an Aurora instrument (RedShift BioAnalytics, Inc.) at 1 mg/mL protein concentration. The samples were analyzed against the matching buffer (50 mM sodium phosphate pH 7.0, 50 mM NaCl, 0.5 mM DTT) at a backing pressure between 7-12 psi with an average flow rate of 1.4 µL/s. The differential absorbance spectra were collected across the Amide-I band from 1588-1712 cm^-1^ with a resolution of 1 cm^-1^.

For thermal ramping measurements, an Aurora TX (RedShift BioAnalytics, Inc.) was used to capture the differential spectral data of sample-buffer over temperature with 0.5 mg/mL CU228 concentration and 1 mM ligand concentration. Equivalent concentrations of ligands were also added to reference buffer for baseline subtraction. Spectra were captured from 1588-1712 cm^−1^ at 1 cm^−1^ resolution. The flow rate was calibrated to 1 μL/min at 25°C and each fluid exchange lasted 0.4 s to remove carryover effects in the MMS cell. Thermal ramping was carried out continuously at 1°C/min from 25 to 75°C. Raw spectra were captured approximately every 17 s along with cell temperature measured by a thermistor attached to the MMS flow cell. Dark scan, sample channel, and reference channel measurements were captured 3 s apart. Due to the continuous temperature ramp, dark scan and reference detector signals were phase-corrected to align them in time and temperature with sample measurements. The sample and reference channel signals were dark offset corrected, and water vapor was subtracted to yield differential absorbance.

The *delta* analytics software (Lucas et al. 2025) was used for sample analysis and spectral processing. The differential absorbance spectra were normalized to absolute spectra using a 0.63 nominal displacement factor using apo-CU228 as the model protein for the fitting. Absolute spectra were then converted to similarity spectra by first taking the second derivative, then inversed and baselined. Savitsky-Golay smoothing was applied to the second derivative plot using a window of 19 wavenumbers. HOS structural elements were calculated using the similarity spectra by Gaussian curve fitting. A list of Gaussians with designated wavenumbers and corresponding structural types is shown in **Supplemental Table 1** (Kendrick et al. 2020; Liu et al. 2020; Ma et al. 2023; Wei et al. 2025).

### Nuclear Magnetic Resonance

^15^N/^1^H TROSY spectra were collected with a Bruker Avance III HD 800 MHz spectrometer with a 5 mm TCI CryoProbe at 298 K using the WADE-TROSY (Manu et al. 2022) pulse sequence, using 165 µM protein (and 1 mM of small molecule ligands, when indicated) in 10 mM sodium phosphate pH 7.0, 50 mM NaCl, and 0.5 mM DTT. Spectra were acquired in 2 hr each. 1D ^1^H NMR spectra were acquired of apo CU228 protein in the absence and presence of 1.25% DMSO at 700 MHz using standard Bruker pulse sequences as well. Data were processed and analyzed using NMRfx Analyst (Norris et al. 2016; Koag et al. 2025).

Saturation transfer difference (STD) spectra were collected with a Bruker Avance III HD 700 MHz spectrometer with a 5 mm QCI-F CryoProbe at 298K using standard Bruker pulse sequences. Samples contained 20 µM CU228 and indicated ligand concentrations of 200, 400, 600, 800, and 1000 µM in the same 10 mM sodium phosphate pH 7.0, 50 mM NaCl, and 0.5 mM DTT buffer as used for other NMR experiments. Saturation transfer difference spectra were collected using the pulse program “stddiffesgp.3” with varying saturation times of 0.5, 0.75, 1, 1.5, 2, 2.5, 3.5, and 5 s. The interscan relaxation delay (d1) was kept constant at 5 s for all the experiments. Binding affinities were calculated as previously reported (Mayer and Meyer 2001; Angulo et al. 2010). To calculate saturation transfer difference (STD) values, absolute integrals of the difference spectra for three separate spectral regions (8.42-8.39 ppm, 7.83-7.78 ppm, and 7.42-7.26 ppm) were calculated and normalized over the reference integral (I_0_). The STD amplification factor (STD-AF) was calculated using:

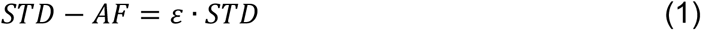

where ε = ([L]_total_/[P]_total_), making STD-AF proportional to the fraction of bound protein. Once plotted against delay time, the data were fit to the STD buildup curve equation:

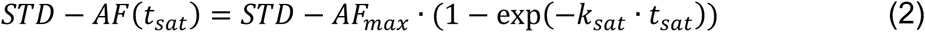

Individual variables can be seen in **Table S3**. STD-AF_0_ was then calculated by multiplying STD-AF_max_ x k_sat._ Plotting STD-AF_0_ versus ligand concentration allowed least-squares fitting to a Langmuir curve:

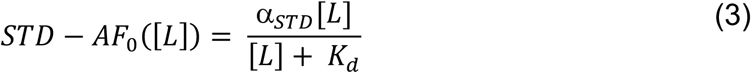

Data processing and absolute integrals were calculated using NMRfx Analyst (Norris et al. 2016; Koag et al. 2025). Curve fitting and analysis were done using R Statistical Software v4.4.2 (Elzhov TV 2023; Wickham H 2023; RCoreTeam 2024).

## Acknowledgements

This work was supported by grants from the NIH (R01 GM106239 and R35 GM156296 to K.H.G.). The authors would like to thank members of the Gardner Laboratory for useful discussions, Dr. Jia Liu (Director, CUNY ASRC Epigenetics Core) for assistance with the QuantStudio 7 Flex Real-time qPCR machine protocol to undertake the thermal shift assay, and Dr. Bruce Johnson (Senior Research Director, Computational Sciences, CUNY ASRC Structural Biology Initiative) for advice on the use of NMRfx Analyst.

## Notes

### Competing Interest Statement

Author R.H.H. is a paid employee of RedShift BioAnalytics, Inc., manufacturer of the Aurora instruments used to collect and interpret MMS data in this manuscript.

### Summary of Updates

Figures 4 and 5 revised to simplify presentation of MMS data and more clearly show concentration-dependent ligand effects Minor text revisions to address grammatical errors

