## Supporting Information for "A Pipeline for Screening Small Molecule-Enhanced Protein Stability in A Bacterial Orphan Receptor"

Contents:

Figures S1-S7

Tables S1-S3

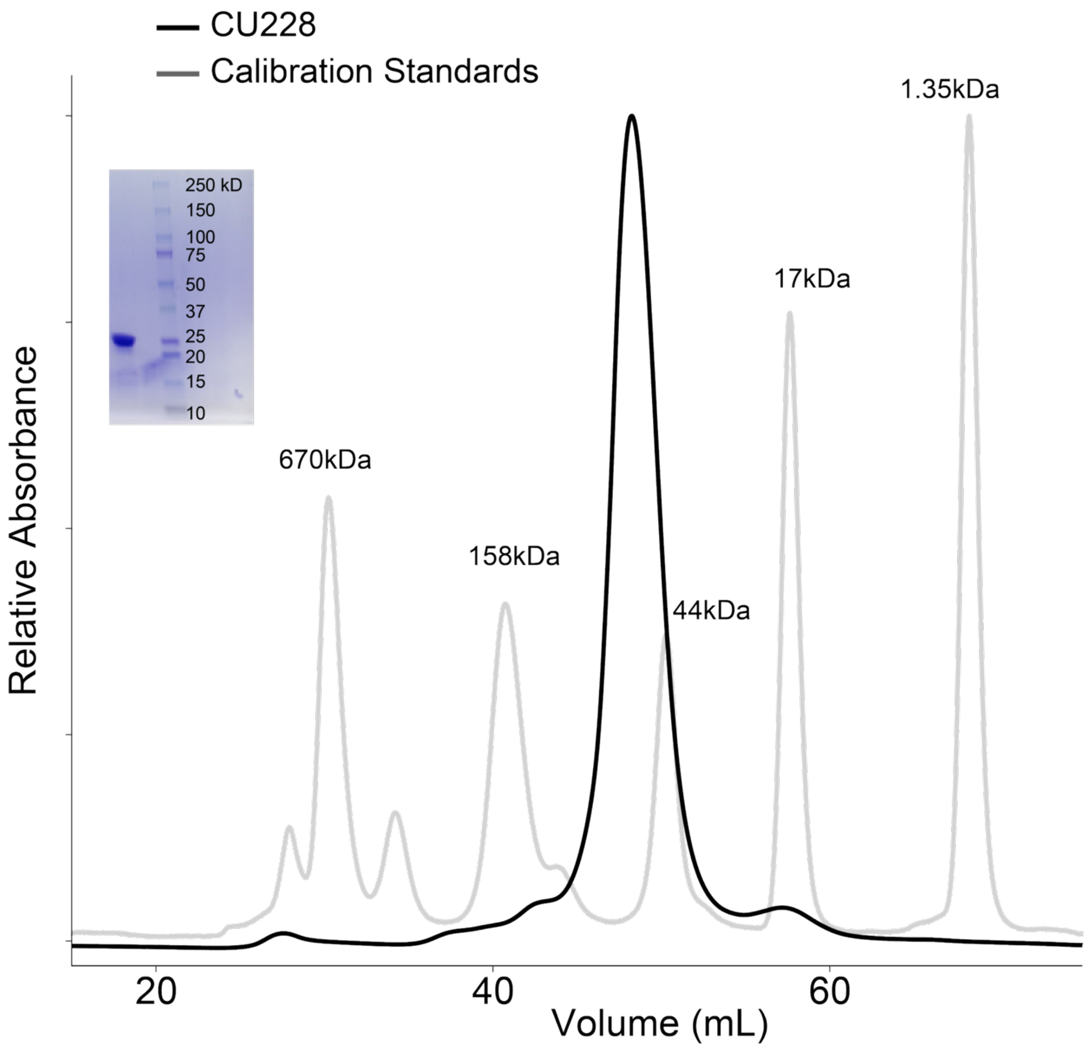


**Figure S1. CU228 purification.** SEC chromatogram of CU228 (black) and chromatography standards (grey). The insert shows SDS-PAGE gel of purified CU228.

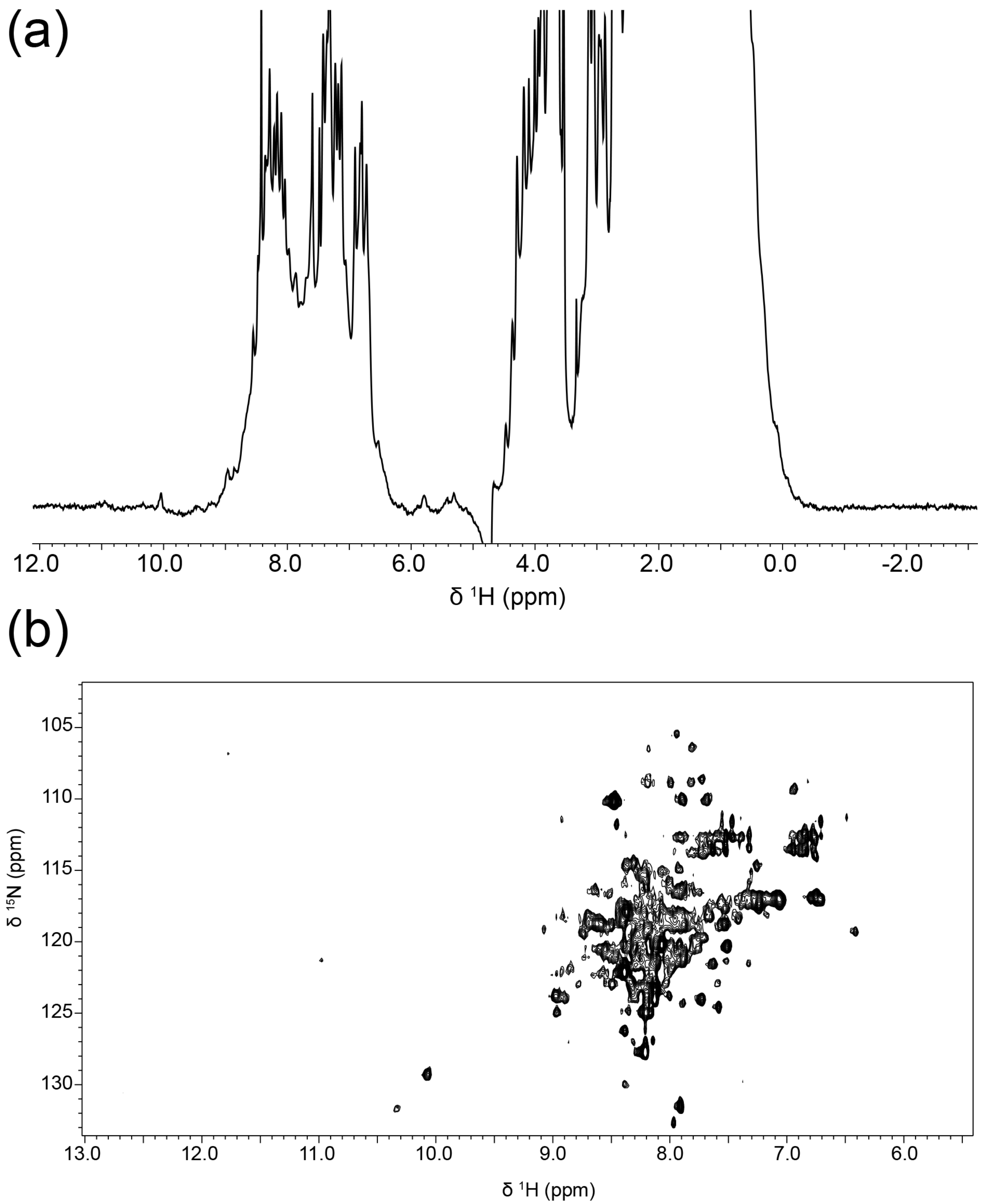


**Figure S2. Initial characterization of apo CU228 by 1D and 2D protein NMR.** (a) $^{1}H$ NMR of CU228. (b) $^{15}N/^{1}H$ TROSY of CU228.

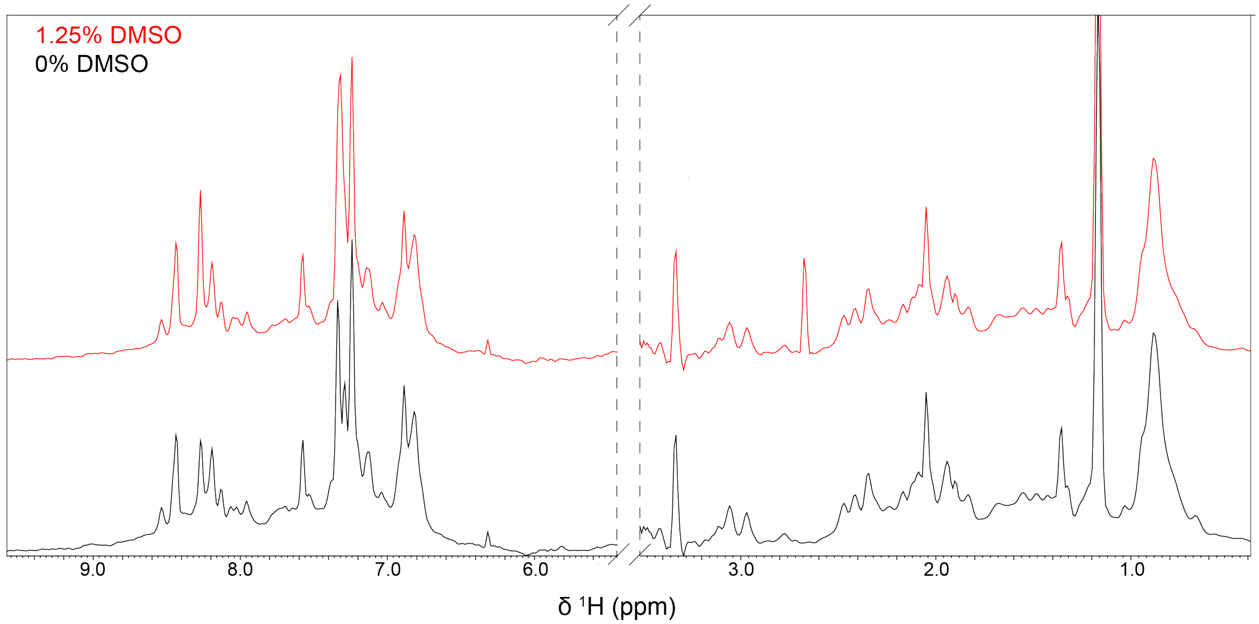


**Figure S3. $^1$H NMR with CU228 in the presence and absence of 1.25% DMSO-d6.** Aliphatic and amide sections of a 1D $^1$H spectra of 136 μM CU228 in the absence (black) or presence (red) of 1.25% DMSO-d6.

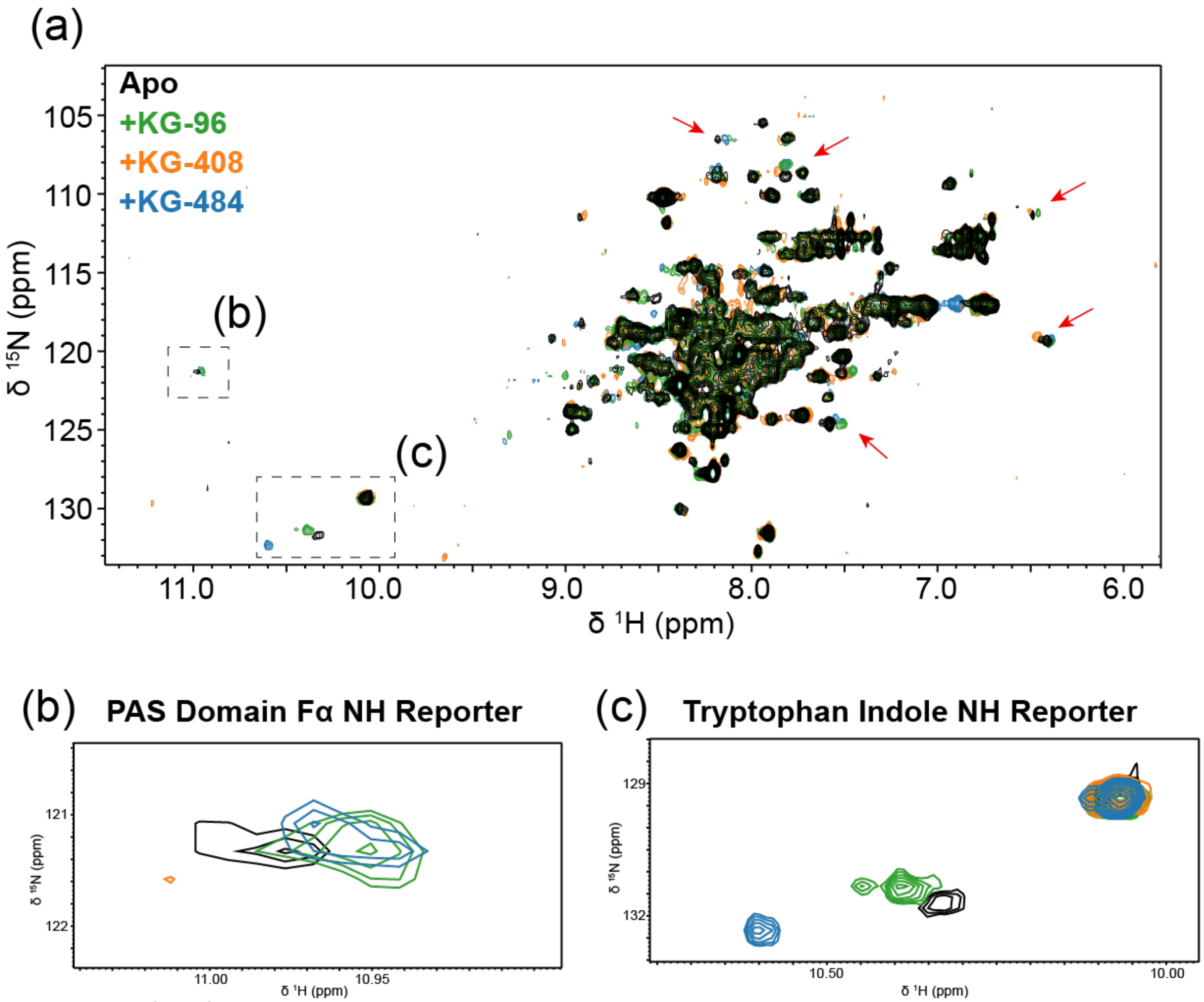


**Figure S4. $^{15}N/^{1}H$ TROSY spectra of CU228 alone and the presence of small molecule ligands.** (a) TROSY spectra of 165 µM CU228 (black) and with the addition of 1 mM KG96 (green), KG408 (orange), or KG484 (blue). Red arrows indicate peaks showing ligand-dependent chemical shift changes, with specific compounds giving rise to specific patterns of changes. (b) Expansion of spectral region associated with a hallmark PAS domain peak arising from a backbone amide at the N-terminus of the Fα helix as detailed in the text. (c) Expansion of spectral region typically occupied by tryptophan sidechain indole NH groups. Four Trp residues are present in CU228, all of which are located in the PAS domain.

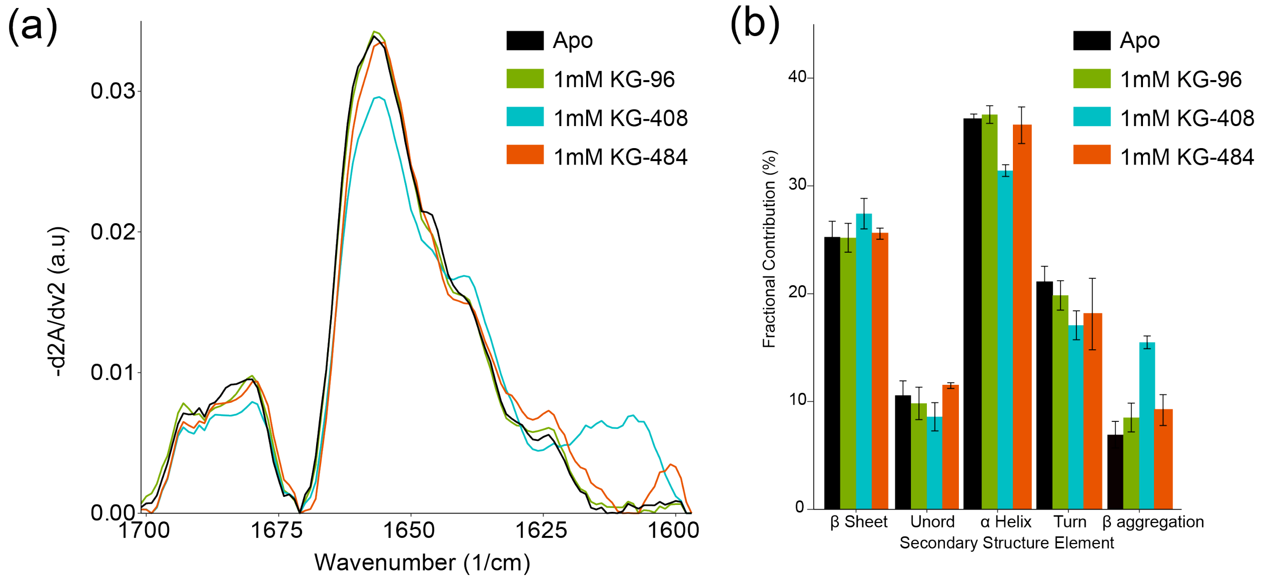


**Figure S5. MMS of CU228 with small molecules at 25°C.** (a) Second derivative Gaussian fits of IR spectra recorded for Apo CU228 and CU228 with 250 μM KG-96, KG-408, and KG-484. (b) Higher-order structure fractional contribution of each sample calculated from traces in panel a.

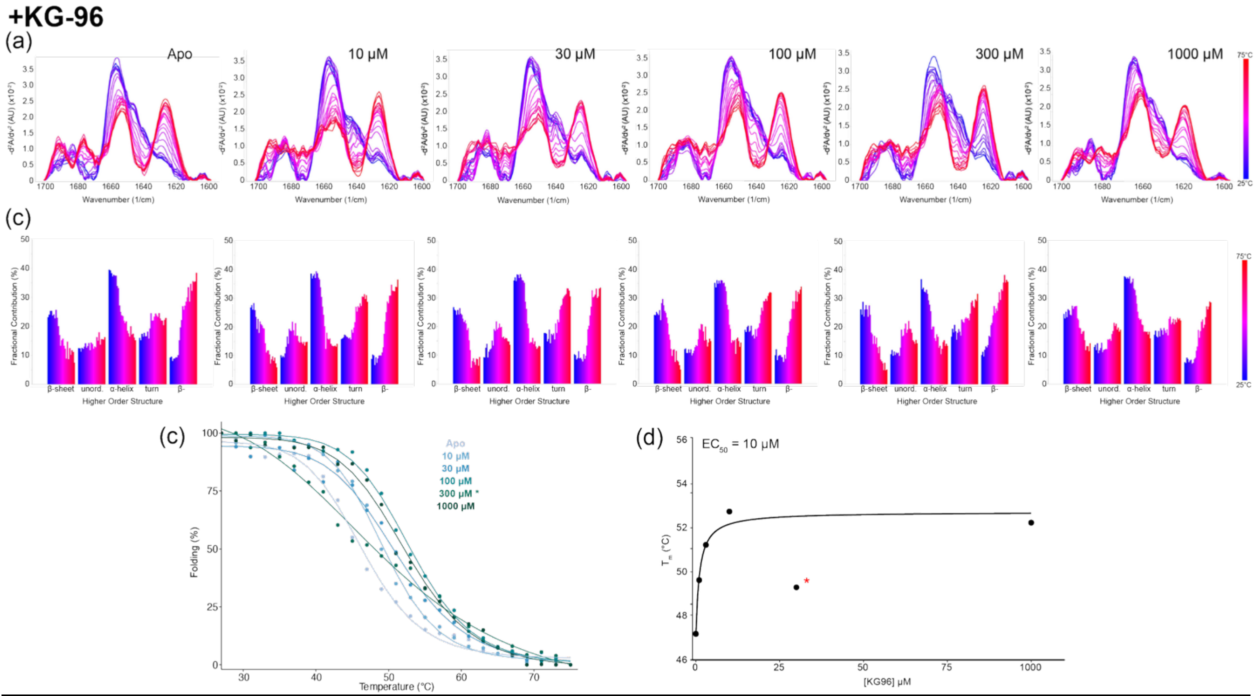


**Figure S6. Concentration Dependent Thermal ramp MMS of CU228 and KG-96.** (a) IR traces across from 25-75°C at five different concentrations. (b) Higher order structure fractional contribution as a function of temperature for each titration point. (c) Global melting curves of CU228 with addition of each KG-96 concentration. (d) KG-96 impact on the melting temperature of CU288 as a function of ligand concentration. Red asterisk indicates that the 300 μM point was omitted from fitting.

**+KG-484**

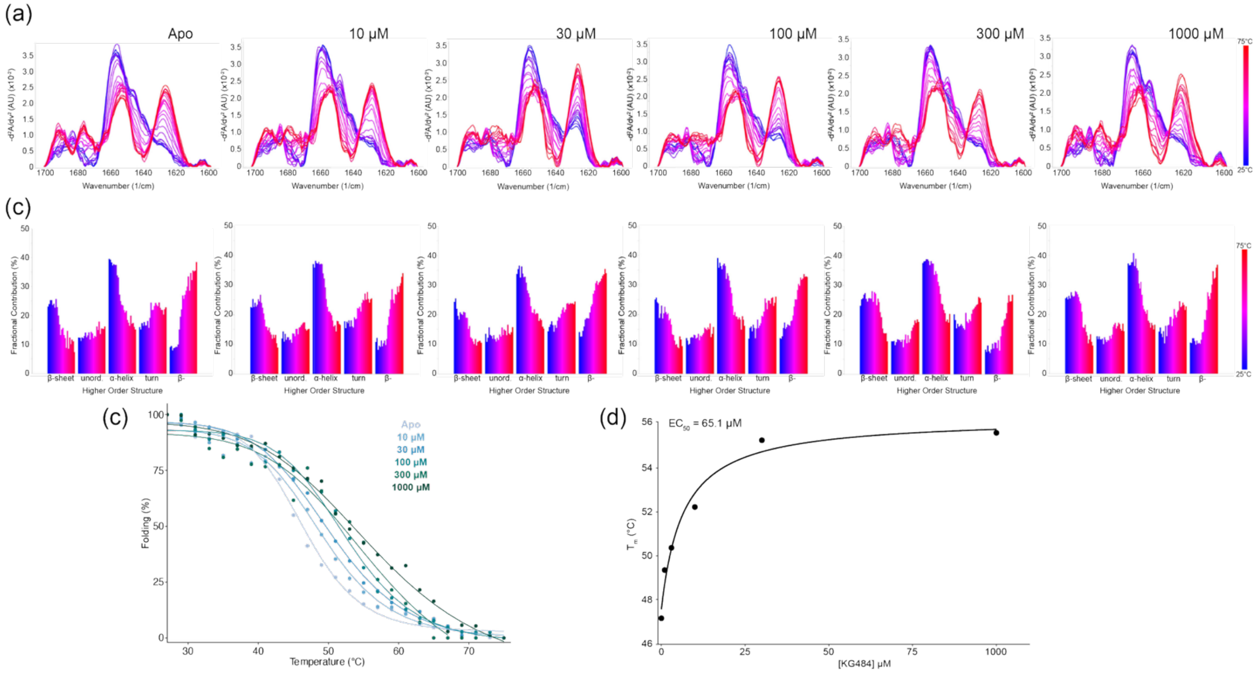


**Figure S7. Concentration Dependent Thermal ramp MMS of CU228 and KG-484.** (a) IR Traces across from 25-75°C at five different concentrations. (b) Higher order structure fractional contribution as a function of temperature for each titration point. (c) Global melting curves of CU228 with addition of each KG-484 concentration. (d) KG-484 impact on the melting temperature of CU288 as a function of ligand concentration.

| Wavenumber ($cm^{-1}$) | Secondary Structure |
|---|---|
| 1618 | Beta- (intermolecular β-sheet) |
| 1624 | Beta- |
| 1627 | Beta- |
| 1632 | Beta |
| 1638 | Beta |
| 1642 | Beta |
| 1650 | Unordered |
| 1656 | Alpha |
| 1666 | Turn |
| 1672 | Turn |
| 1680 | Turn |
| 1688 | Turn |
| 1695 | Beta- |

**Table S1.** Peak assignments for Gaussian curve fitting.

| Ligand | [Ligand] (μM) | $T_m$ (β-) | $T_m$ (β-sheet) | $T_m$ (α-helix) |
|---|---|---|---|---|
| **Apo** | - | 48.3 | 47.0 | 43.3 |
| **KG-96** | 10 | 49.0 | 49.6 | 47.2 |
| | 30 | 51.3 | 52.3 | 47.2 |
| | 100 | 51.8 | 54.6 | 47.3 |
| | 300 | 42.3 | 53.9 | 44.1 |
| | 1000 | 53.4 | 60.0 | 47.8 |
| **KG-408** | 10 | 48.6 | 49.2 | 44.3 |
| | 30 | 51.9 | 51.8 | 45.7 |
| | 100 | 51.7 | 53.3 | 45.1 |
| | 300 | 53.7 | 55.4 | 45.0 |
| | 1000 | 54.4 | 60.6 | 46.5 |
| **KG-484** | 10 | 50.5 | 51.2 | 46.7 |
| | 30 | 49.4 | 52.0 | 47.8 |
| | 100 | 53.4 | 55.7 | 51.1 |
| | 300 | 55.7 | 60.0 | 52.5 |
| | 1000 | 53.3 | 61.0 | 47.5 |

**Table S2: Secondary structure melting temperatures ($T_m$, in °C) of CU228 with and without small molecule ligands.** KG-96, KG-408, and KG-484 were added at five different concentrations as indicated. (β- = intermolecular β-sheet aggregation)

| Ligand | Peak Region | [Ligand] (μM) | STD-$AF_{max}$ | $k_{sat}$ ($s^{-1}$) | STD-$AF_0$ (STD-$AF_{max}$ x $k_{sat}$) |
|---|---|---|---|---|---|
| KG96 | 1 | 0 | - | - | 0 |
| | | 200 | 1.16 | 0.0632 | 0.073312 |
| | | 400 | 0.744 | 0.123 | 0.091512 |
| | | 600 | 1.39 | 0.066 | 0.09174 |
| | | 800 | 1.22 | 0.0763 | 0.093086 |
| | | 1000 | 0.52 | 0.232 | 0.12064 |
| | 2 | 0 | - | - | 0 |
| | | 200 | 1.75 | 0.0578 | 0.10115 |
| | | 400 | 1.26 | 0.11 | 0.1386 |
| | | 600 | 1.9 | 0.0882 | 0.16758 |
| | | 800 | 1.26 | 0.142 | 0.17892 |
| | | 1000 | 1.28 | 0.156 | 0.19968 |
| | 3 | 0 | - | - | 0 |
| | | 200 | 0.912 | 0.212 | 0.193344 |
| | | 400 | 1.08 | 0.187 | 0.20196 |
| | | 600 | 1.65 | 0.129 | 0.21285 |
| | | 800 | 1.16 | 0.207 | 0.24012 |
| | | 1000 | 1.18 | 0.21 | 0.2478 |
| KG408 | 1 | 0 | - | - | 0 |
| | | 200 | 1.21 | 0.513 | 0.62073 |
| | | 400 | 2.33 | 0.426 | 0.99258 |
| | | 600 | 3.52 | 0.336 | 1.18272 |
| | | 800 | 4.09 | 0.296 | 1.21064 |
| | | 1000 | 4.73 | 0.327 | 1.54671 |
| | 2 | 0 | - | - | 0 |
| | | 200 | 1.57 | 0.505 | 0.79285 |
| | | 400 | 3.03 | 0.41 | 1.2423 |
| | | 600 | 3.83 | 0.367 | 1.40561 |
| | | 800 | 4.41 | 0.362 | 1.59642 |
| | | 1000 | 5.44 | 0.302 | 1.64288 |
| | 3 | 0 | - | - | 0 |
| | | 200 | 1.36 | 0.508 | 0.69088 |
| | | 400 | 2.61 | 0.43 | 1.1223 |
| | | 600 | 3.57 | 0.389 | 1.38873 |
| | | 800 | 4.39 | 0.328 | 1.43992 |
| | | 1000 | 5.27 | 0.313 | 1.64951 |
| KG484 | 1 | 0 | - | - | 0 |
| | | 200 | 0.0747 | 0.821 | 0.0613287 |
| | | 400 | 0.326 | 0.113 | 0.036838 |
| | | 600 | 0.123 | 0.844 | 0.103812 |
| | | 800 | 0.259 | 0.308 | 0.079772 |

| | | | | | |
|---|---|---|---|---|---|
| | | 1000 | 0.185 | 0.62 | 0.1147 |
| | 2 | 0 | - | - | 0 |
| | | 200 | 9.48 | 0.00214 | 0.0202872 |
| | | 400 | 0.246 | 0.16 | 0.03936 |
| | | 600 | 1.21 | 0.0274 | 0.033154 |
| | | 800 | 1.76 | 0.0254 | 0.044704 |
| | | 1000 | 0.533 | 0.111 | 0.059163 |
| | 3 | 0 | - | - | 0 |
| | | 200 | 0.0552 | 0.276 | 0.0152352 |
| | | 400 | 11.9 | 0.00178 | 0.021182 |
| | | 600 | 0.178 | 0.155 | 0.02759 |
| | | 800 | 6.91 | 0.00418 | 0.0288838 |
| | | 1000 | 0.276 | 0.152 | 0.041952 |

**Table S3:** Variables derived from STD buildup curves (**Eq. 2**) used to plot Langmuir fittings.
